# Genetic Association of The CYP7A1 Gene with Duck Lipid Traits

**DOI:** 10.1101/456475

**Authors:** Yuan-Yu Qin, Yi-Yu Zhang, Hua-Lun Luo, Lei Wu

## Abstract

Cholesterol 7α-hydroxylase (Cyp7a1) participates in lipid metabolism of liver, and its pathway involves catabolism of cholesterol to bile acids and excretion from the body. However, little is known about the effect of the polymorphisms of CYP7A1 gene on duck lipid traits. In the present study, seven novel synonymous mutations loci in exon 2 and exon 3 of CYP7A1 gene in Cherry Valley ducks were identified using PCR production direct sequencing. One novel SNP g.1033130 C>T was predicted in exon 2. Six novel SNPs g.1034076 C>T, g.1034334 G>A, g.1034373 G>A, g.1034448 T>C, g.1034541 C>G, and g.1034550 G>A were discovered in exon 3. Six haplotypes were detected using SHEsis online analysis software, and five loci (g.1034334G>A, g.1034373G>A, g.1034448T>C, g.1034541C>G, and g.1034550G>A) were in complete linkage disequilibrium, and as a block named Locus C3. By single SNP association analysis, we found that the g.1033130 C>T locus was significantly associated with IMF, AFP, TG, and TC (*P*<0.01 or *P*<0.05) respectively, the g.1034076 C>T locus was significantly associated with AFP (P<0.05), and the locus C3 was significantly associated with TCH (*P*<0.05). Sixteen dipoltypes were detected by the combination of haplotypes, and demonstrated strong association with IMF, AFP, TG, and TCH (*P*<0.01). Therefore, our data suggested that the seven SNPs of CYP7A1 gene are potential markers for lipid homeostasis, and may be used for early breeding and selection of duck.

---

Lipid metabolism is a dynamic biological process that includes lipid synthesis, accumulation, distribution to different tissues, and excretion procedure. The molecular mechanism of lipid homeostasis is regulated by a set of well-characterized transport proteins and enzymes that are variously triggered depending on the organismal requirements (Zeng *et al*. 2018). Many signaling pathways involved in lipid metabolism like LXR, PPAR, and SREBP signals are dysregulated, resulting in modification of circulating lipid levels (Chen 2015; Pan *et al*. 2018). In human, the disorder of lipid metabolism leads to many diseases associated with lipid transport, such as type 2 diabetes, fatty liver, and hyperlipidemia and so on (Bjornstad and Eckel 2018). However, the excessive fatty tissue deposition is one of the critical subjects confronted in avian species. Therefore, decreasing body fat deposition and increasing intramuscular fat are main breeding projects in modern poultry industry because consumers change the concept of consumption (Wang *et al*. 2017). But the genetic control of lipid metabolism in poultry is still unclear, and resulting in slow progress for reducing body fatty (Ye *et al*. 2016). The combination of modern molecular biology methods and traditional breeding technologies is the essence of improving breeding efficiency in the future.

Cholesterol 7α-hydroxylase (CYP7A1), which is an important rate-limiting enzyme in bile acid synthetic pathway in human and animal liver and therefore plays a crucial function in maintaining cholesterol homeostasis (Wang *et al*. 2018). The promoter of CYP7A1 gene includes deeply conservative bile acid responsive regions known to be regulated by feedback repression by different transcription factors in reaction to rising the concentration of hepatic bile acid (Chiang 2009). In human, the negative feedback mechanism of CYP7A1 regulation is the small heterodimer partner and farnesoid X receptor regulatory cascade (Lin 2015). For example, a bile acid chenodeoxycholic acid activates farnesoid X receptor to increase transcription of small heterodimer partner, and inhabits CYP7A1 activation by repression of transcription factors, hepatocyte nuclear factor 4α and liver receptor homolog-1, which are indispensable for CYP7A1 transcription (Al-Aqil *et al*. 2018). In addition, the negative feedback mechanism on CYP7A1 is discovered in human/rodent intestine, where activation of farnesoid X receptor induces a hormonal signaling molecule fibroblast growth factor 15/19 that suppresses CYP7A1 via interaction with the liver fibroblast growth factor receptor 4 through the c-Jun signaling pathway (Memon *et al*. 2018). Liver X receptor α (LXRα), an oxysterol-binding transcription factor, which regulate the transcription level of CYP7A1 (Yang *et al*. 2018). CYP7A1 was downregulated with lithogenic time progressed, while CYP7A1 mRNA was increased 3-fold under the treatment of Yinchenhao Decoction, and revealed that CYP7A1 mRNA may be efficient targets of Yinchenhao Decoction, which may be a preventive medicine against cholesterol gallstone formation (Meng *et al*. 2018). CYP7A1 mediate the transformation of cholesterol into bile acids in vitro. In Farnesoid X receptor (Fxr) ^-/-^ and wild-type mice with hypercholesterolemia, injection of 1,25(OH)_2_D_3_ consistently reduced levels of liver and plasma cholesterol, and increased the levels of hepatic CYP7A1 mRNA and protein; in mouse and human hepatocytes, incubation with 1,25(OH)_2_D_3_ increased the CYP7A1 mRNA level, and reduced cholesterol (Chow *et al*. 2014). In neonatal piglets, formula feeding resulted in fecal bile acid loss and increased CYP7A1 protein expression, which is associated with decreased effectivity in inhibiting CYP7A1 expression via SHE and FGF 19 transcriptional repression mechanisms (Mercer *et al*. 2018). The latest research showed that some miRNAs like miR-17, miR-34a, miR-122, are the regulator of CYP7A1 signaling in hepatic lipid metabolism, suggesting a possible improvement pathway for lipid deposition (Gong *et al*. 2018; Cui *et al*. 2018; Zinkhan *et al*. 2018).

Based on previous evidence, CYP7A1 plays a notable role in human and animal lipid metabolism, and is an important candidate gene. The intent of this study detected whether the polymorphisms of CYP7A1 gene are associated with duck lipid traits. To test this hypothesis, we genotyped 7 novel single nucleotide polymorphisms (SNPs) of CYP7A1 gene in blood samples from Cherry Valley ducks and identified these SNPs that are related to lipid traits.

## MATERIALS AND METHODS

### Experimental Animals

A total of 205 females of Cherry Valley ducks which is an introduced duck variety were used in present study. All of individuals were hatched on the same day, and raised in a semi-open house and fed commercial corn–soybean diets based on the NRC requirements and subjected to same management conditions in poultry research institute of Guizhou University, Guiyang, Guizhou, P.R.China. All of ducks were euthanized that the specific implementation plan was taken all experimental ducks placed in an operating room filled with a mixture of 90% argon and 10% nitrogen at 70 days of age, and then the ducks were quickly bled, and collected blood samples and dissected after they were unconscious. All experiment procedures were performed according to the Laboratory animal—Guideline for ethical review of animal welfare of China (permit number: GB/T 35892-2018) that was issued by China Laboratory Animal Standardization Technology Committee (SAC/TC 281).

### Meat Quality Measurement

Chest muscle samples were rapidly collected and stored at –20°C. Intramuscular fat (IMF) content of chest muscles and percentage of abdominal fat (AFP) of 205 females of Cherry Valley ducks was measured using the Soxhlet extraction method (Li *et al*. 2018) and slaughter segmentation method (Lin *et al*. 2018), respectively. Serum was isolated at 3,500 rpm, 4°C for 10 min from blood that was held at room temperature for 1 h, and then stored at –30°C. The contents of triglycerides (TG) and total cholesterol (TCH) in serum were detected using qualified laboratory methods (He *et al*. 2018).

### Genomic DNA Extraction

Genomic DNA was extracted from whole blood samples according to the manufacturer’s instructions of QIAamp DSP DNA Blood Mini Kit (Shanghai Labpal Co. Ltd., China). The quantity and quality of genomic DNA was measured by ND-8000 spectrophotometer (Thermo Fisher Scientific) and 1% agarose gel electrophoresis respectively, and stored at −20°C.

### Primer Design and Polymerase Chain Reaction Conditions

Four pairs of primers were designed using Primer 3 Input (version 0.4.0) (http://bioinfo.ut.ee/primer3-0.4.0/) based on the *Anas platyrhynchos* reference genomic sequence (GenBank accession: NW_004676497.1) and duck CYP7A1 gene mRNA sequence. The primers were synthesized by Invitrogen Co. Ltd. (Peking, China) (Table 1). Polymerase chain reaction (PCR) was performed in a total volume of 25 µL consisted of 10 μL 2×Pfu Master Mix (Peking Cwbiotech Co. Ltd., China), 1 µL each of upstream and downstream primer (10 µmol/L), 1μL of template genomic DNA (100ng/μL), and 12 µL of ddH_2_O. The PCR protocol was performed as follows: preliminary denaturation denaturation at 94°C for 8 min, followed by 35 cycles of denaturing at 94°C for 40 s, annealing at for 45 s at the optimum primer annealing temperature, elongation for 45 s at 72°C, and a final extension for 10 min at 72°C. PCR products were stored at 4°C.

**Table 1.**
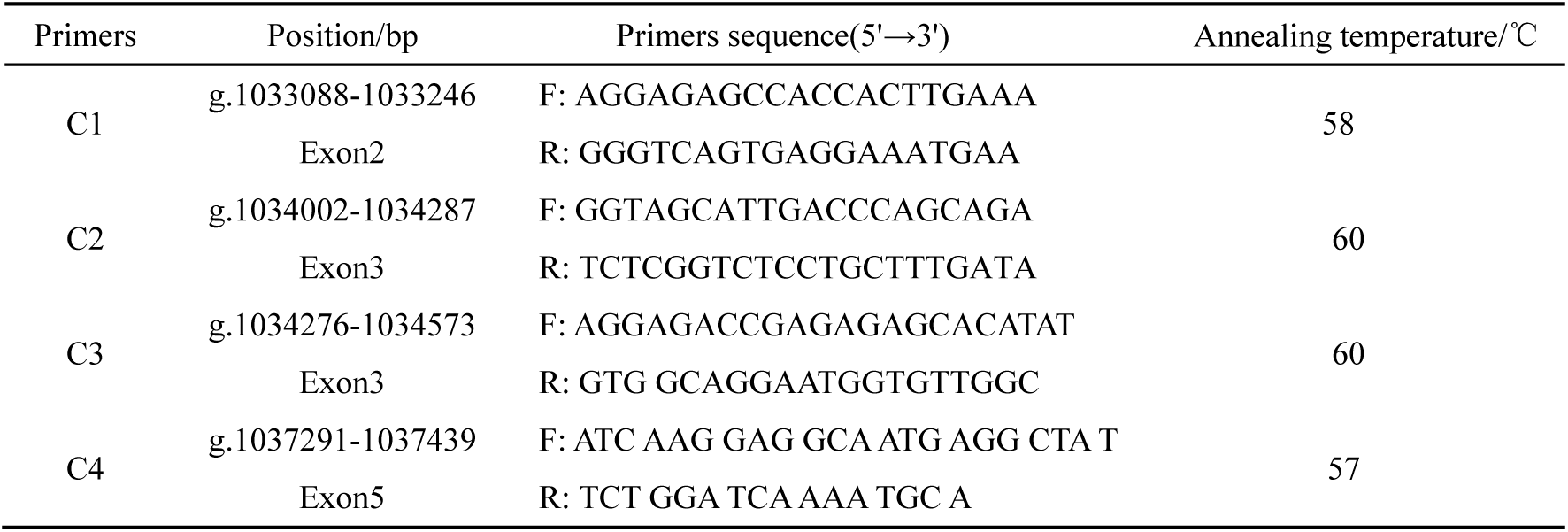
Primers information used for PCR amplification of duck CYP7A1 gene.

### Gene Polymorphism Analysis

A 5µL of each PCR product was detected by 1.5% agarose gel electrophoresis, visualized with gold view staining, and quantified using Invitrogen®E-Gel Imager (Thermo Fisher Scientific). The remaining PCR products were purified according to the manufacturer’s instructions of SanPrep Column PCR Product Purification Kit (Sangon Biotech Co. Shanghai, China), and then were directly sequenced using HiSeq 3000/HiSeq 400 analyser (Illumina, Inc. USA). The sequencing results were analysed with AlignIR™ Software (LI-COR Corporate, Nebraska, USA) to determine the single nucleotide polymorphisms (SNPs) of CYP7A1 in Cherry Valley ducks.

### Statistical Analysis

D'/r2 value of pair-loci linkage disequilibrium (LD), haplotype frequency, diplotype frequency, genotype frequency, allele frequency, and chi-square (*χ2*) of Hardy-Weinberg equilibrium tests for each locus were calculated using SHEsis platform (http://analysis.bio-x.cn/). Effective number of alleles (Ne), heterozygosities (He), and polymorphism information content (PIC) were calculated using Cervus 3.0 software. Statistical analyses were performed using SPSS 16.0 (SPSS Inc., Chicago, IL, USA). The linear model was used: Y=µ+ G + e, Y - lipid trait, G - the genotype effect or diplotype effect, µ - the mean for each trait, e - the random error. Statistical significance of the difference for each lipid trait between genotypes and diplotypes was evaluated using Post Hoc multiple comparisons for observed means. Data were showed as mean ± SE.

## RESULTS

### SNPs Identification of CYP7A1 Gene

Four pairs of different primers for CYP7A1 gene in Cherry Valley ducks were detected for SNPs. Through the sequence alignment of 205 samples, no polymorphic site was found in primer C4. One novel SNP g.1033130 C>T was predicted in exon 2. Six novel SNPs g.1034076 C>T, g.1034334 G>A, g.1034373 G>A, g.1034448 T>C, g.1034541 C>G, and g.1034550 G>A were discovered in exon 3. The aligning maps showing different genotypes were displayed in Fig. 1. Seven novel SNPs in exon 2 and exon 3 have no caused the amino acids change, which belonged to synonymous mutations, and produced three genotypes of each SNP.

**Figure 1.**
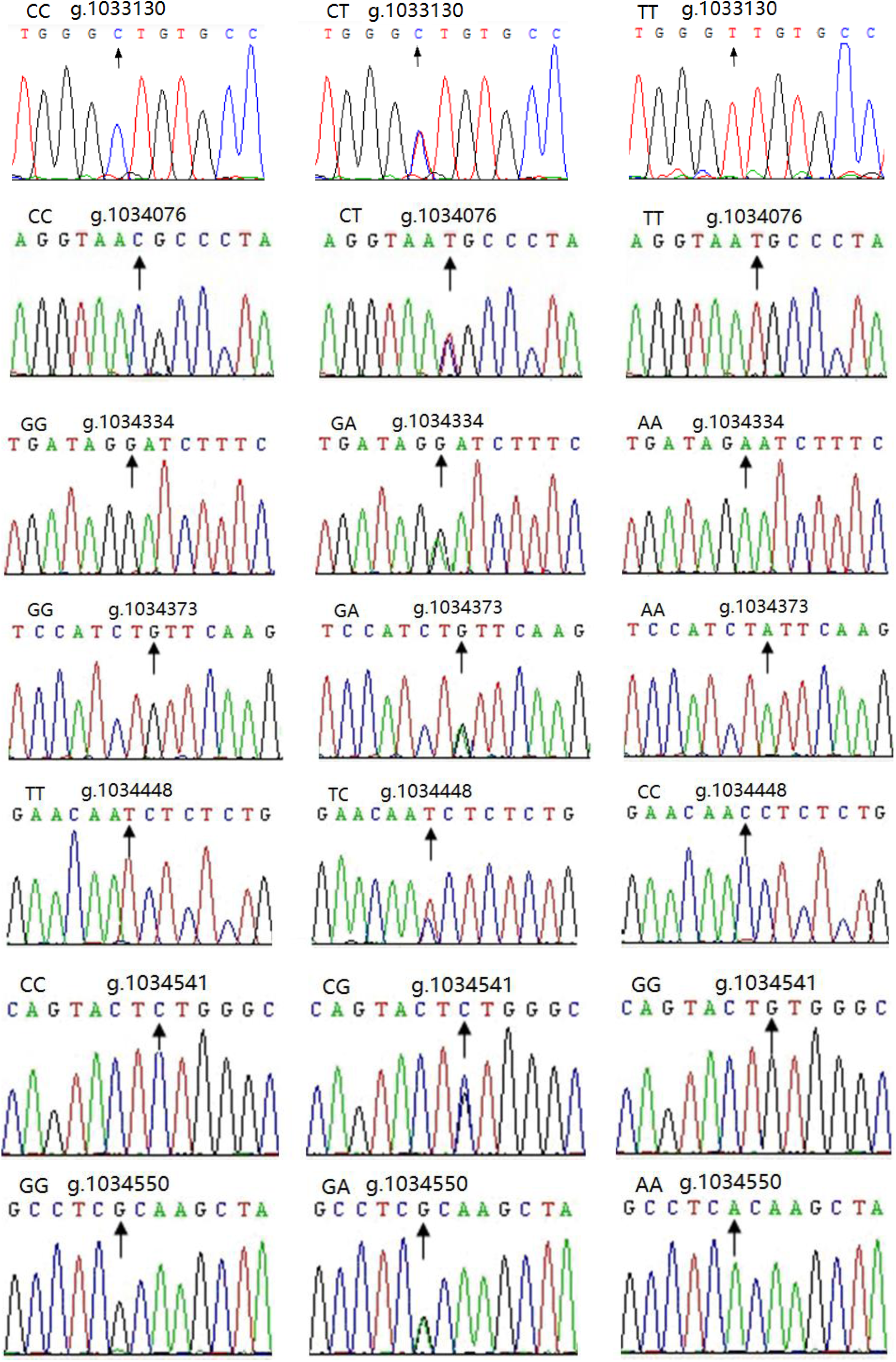
Sequencing results for novelly discovered SNP sites

### Linkage Disequilibrium, Haplotypes and Diplotypes Analysis of Seven SNPs of CYP7A1 Gene

Linkage disequilibrium (LD) coefficient D' and r2 between seven SNPs of CYP7A1 gene in Cherry Valley ducks was estimated, and displayed in Table 2 and Figure 2. All of the D' and r2 values between five SNPs g.1034334G>A, g.1034373G>A, g.1034448T>C, g.1034541C>G, and g.1034550G>A were 1, and revealed that they were in complete linkage disequilibrium. Therefore, the five SNPs were taken as a whole for further analysis, and named Locus C3. Subsequent analysis showed that the D' and r2-value between g.1033130C>T and g.1034076 C>T was 0.156 and 0.011 respectively; the D' and r2-value between g.1033130C>T and Locus C3 was 0.192 and 0.019 respectively; furthermore, the D' and r2-value between g.1034076 C>T and Locus C3 was 0.472 and 0.195 respectively. According to the rule that r2>0.33 and |D'|>0.8 are considered to be strong linkage disequilibrium between two alleles (Sharma *et al*. 2018; Ardlie *et al*. 2002; Slatkin 2008), there were no strong linkage disequilibrium between g.1033130C>T, g.1034076 C>T, and Locus C3, respectively.

**Table 2.** Linkage disequilibrium coefficient *D'* and *r^2^* between different SNPs.

| SNPs | g.1033130<br>C>T | g.1034076<br>C>T | g.1034334<br>G>C | g.1034373<br>G>A | g.1034448<br>T>C | g.1034541<br>C>G | g.1034550<br>G>A |
| --- | --- | --- | --- | --- | --- | --- | --- |
| g.1033130C>T | -- | 0.156 | 0.192 | 0.192 | 0.192 | 0.192 | 0.192 |
| g.1034076C>T | 0.011 | -- | 0.472 | 0.472 | 0.472 | 0.472 | 0.472 |
| g.1034334G>A | 0.019 | 0.195 | -- | 1.000 | 1.000 | 1.000 | 1.000 |
| g.1034373G>A | 0.019 | 0.195 | 1.000 | -- | 1.000 | 1.000 | 1.000 |
| g.1034448T>C | 0.019 | 0.195 | 1.000 | 1.000 | -- | 1.000 | 1.000 |
| g.1034541C>G | 0.019 | 0.195 | 1.000 | 1.000 | 1.000 | -- | 1.000 |
| g.1034550G>A | 0.019 | 0.195 | 1.000 | 1.000 | 1.000 | 1.000 | -- |

Note: Upper triangle and down triangle showed Linkage disequilibrium coefficient $D'$ and $r^2$ between different SNPs, respectively.

**Figure 2.**
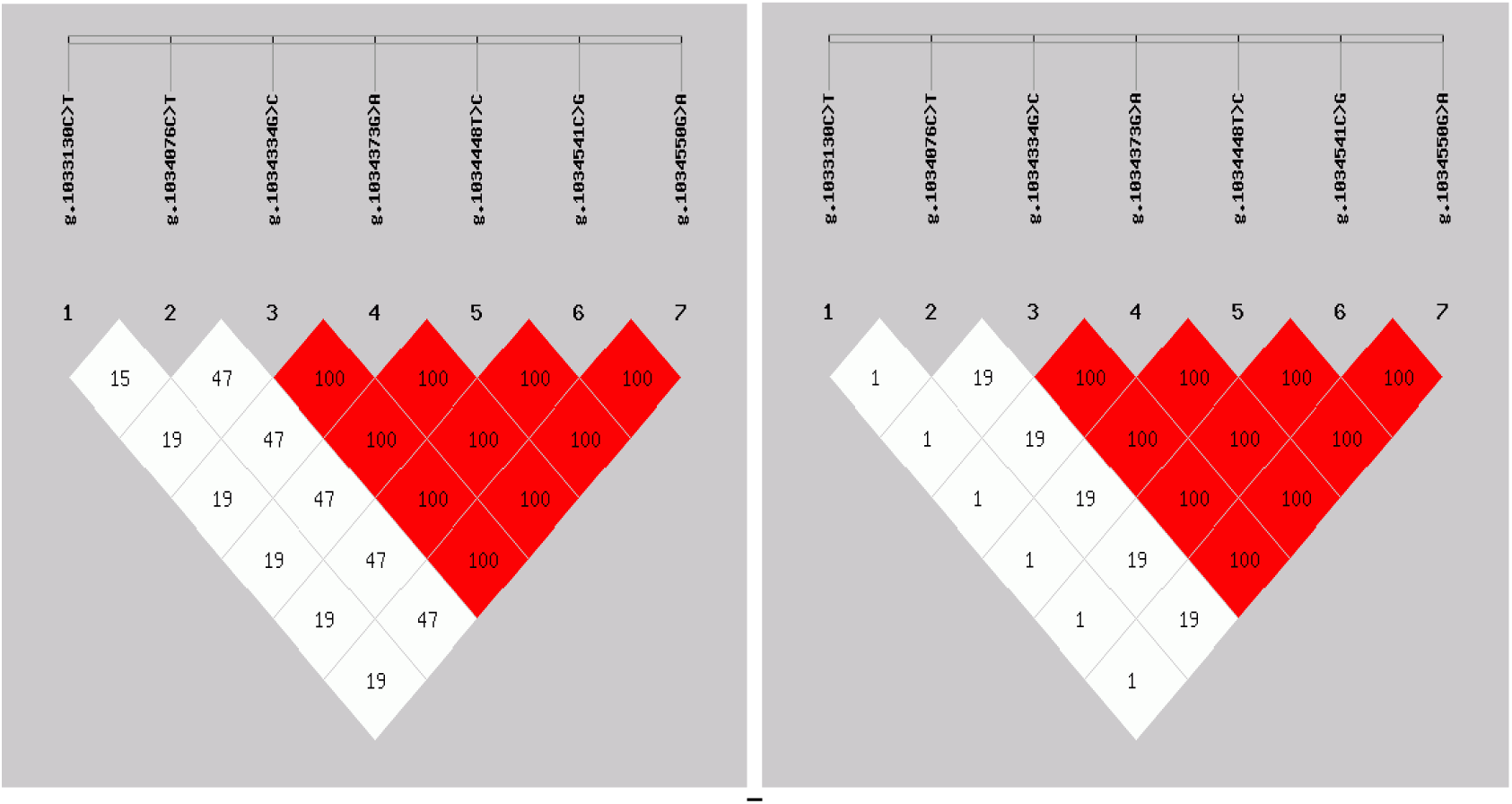
Linkage disequilibrium coefficient between different SNPs (right: r2; left: D')

The haplotypes and diplotypes of seven SNPs of CYP7A1 gene were conducted in Cherry Valley ducks, and showed in Table 3. Six haplotypes were observed in the CYP7A1 gene with one dominant haplotype H1 (CCGGTCG) occurring with a frequency of 0.370, and the frequency of haplotype H6 (TTAACGA) was low (0.100). Totally sixteen diplotypes were found by haplotype combination with two predominant diplotypes H1H1 (CCCCGGGGTTCCGG) and H2H3 (CCCTGAGATCCGGA) with frequency of 0.215 and 0.200 respectively. The frequency of both H5H6 (TTCTGAGATCCGGA) and H6H6 (TTTTAAAACCGGAA) diplotypes was the lowest (only 0.029).

**Table 3.**
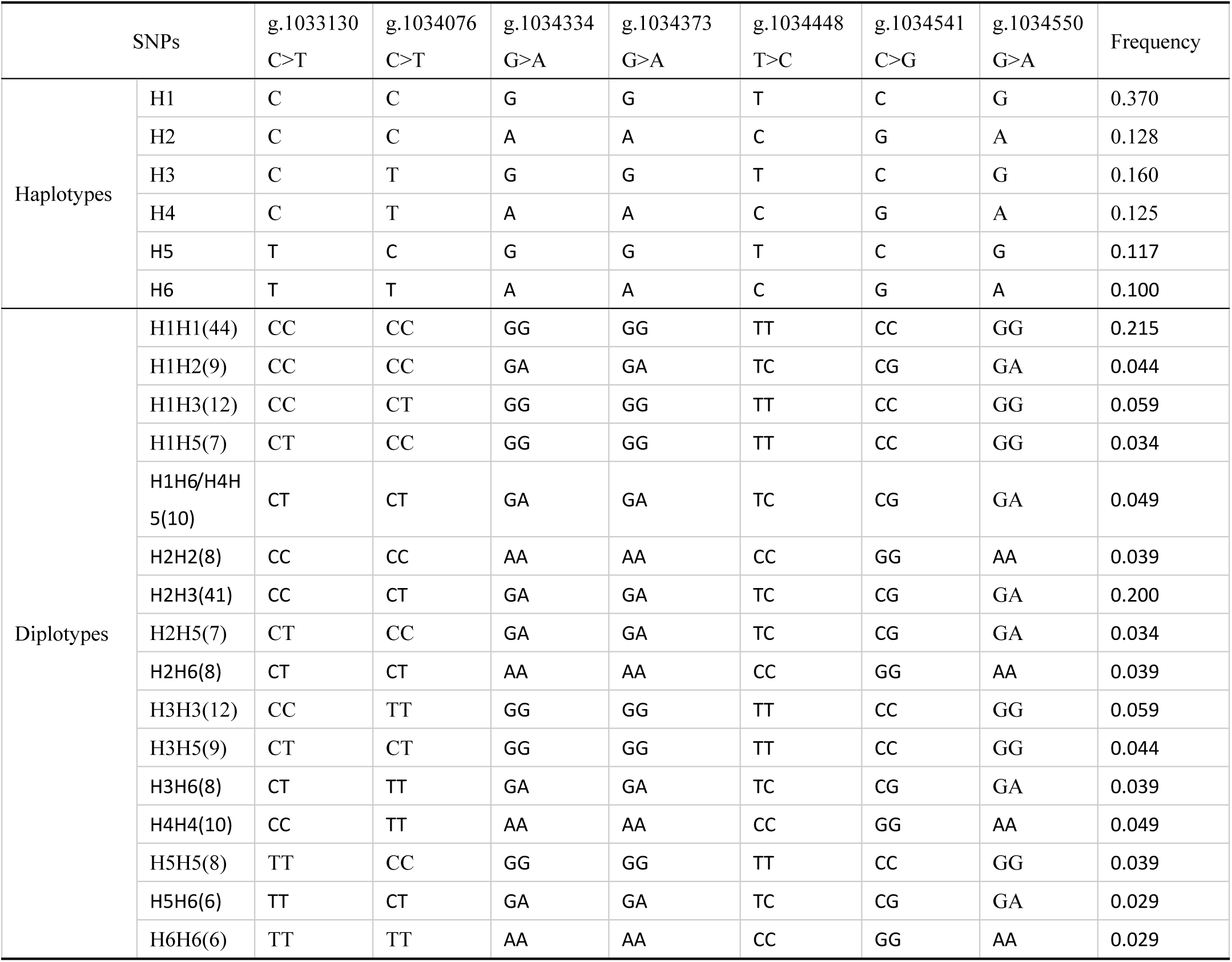
Haplotypes and diplotypes analysis based on the seven SNPs of CYP7A1 gene.

### Genotypic Frequencies, Allelic Frequencies, and Diversity Parameter

The genotype and allele frequency of each SNP in Cherry Valley ducks was summarized in Table 4. Three genotypes, namely, CC, TT, and CT, were detected at the g.1033130 C>T and g.1034076 C>T respectively; AA, AB, and BB were detected at the Locus C3. The frequency of the C, C and A allele of g.1033130 C>T, g.1034076 C>T, and Locus C3 was 0.783, 0.615, and 0.646, respectively, and then the CC, CT and AA genotype was dominant genotype, respectively. The χ2-test indicated that the genotype distribution of g.1033130 C>T locus presents significant deviation from Hardy-Weinberg equilibrium (*HWE*) (*P*<0.01), but the results for the g.1034076 C>T locus and Locus C3 were consistent with HWE (*P*>0.05). The calculated resulted of polymorphism information content (*PIC*) indicated that seven SNPs loci were moderately polymorphic (0.25<*PIC*<0.5).

**Table 4.** Genotype frequency, allele frequency and population genetic information.

| SNPs | Genotype frequency | | | Allele frequency | | *He* | *Ne* | *PIC* | $\chi^2$ |
| --- | --- | --- | --- | --- | --- | --- | --- | --- | --- |
| g.1033130<br>C>T | CC(136) | CT(49) | TT(20) | C | T | 0.340 | 1.515 | 0.282 | 18.057** |
|  | 0.663 | 0.239 | 0.098 | 0.783 | 0.217 |  |  |  |  |
| g.1034076<br>C>T | CC(83) | CT(86) | TT(36) | C | T | 0.474 | 1.900 | 0.361 | 2.684 |
|  | 0.405 | 0.420 | 0.176 | 0.615 | 0.385 |  |  |  |  |
| Locus C3 | AA(92) | AB(81) | BB(32) | A | B | 0.457 | 1.843 | 0.353 | 3.776 |
|  | 0.449 | 0.395 | 0.156 | 0.646 | 0.354 |  |  |  |  |

Note: *He*, heterozygosity; *Ne*, Effective allele number; *PIC*, polymorphism information content; $\chi^2$ -test, Hardy-Weinberg equilibrium, $\chi^2_{0.01(2)}=9.21$ , $\chi^2_{0.05(2)}=5.99$ , \* $P<0.05$ , \*\* $P<0.01$ .

### Associations between Identified SNPs and Four Lipid Traits

The results of the association analysis between each SNP of CYP7A1 gene and lipid traits in Cherry Valley ducks were shown in Table 5. We found that the g.1033130 C>T locus was significantly associated with IMF, AFP, TG, and TCH respectively (*P*<0.01 or *P*<0.05), the g.1034076 C>T locus was significantly associated with AFP (*P*<0.05), and the locus C3 was significantly associated with TCH (*P*<0.05). At g.1033130 C>T locus, individuals with CC genotype had higher IMF and AFP than those with TT genotype (*P*<0.01), individuals with CC genotype had lower TCH and TG that those with TT genotype (*P*<0.01 and *P*<0.05, respectively), individuals with CT genotype had higher IMF than those with TT genotype (*P*<0.05), and individuals with CT genotype had lower TCH than those with TT genotype (*P*<0.01); at g.1034076 C>T locus, individuals with CC genotype had higher AFP than those with TT genotype (*P*<0.05); at Locus C3, individuals with BB genotype had lower TCH than those with AA genotype (*P*<0.05).

**Table 5.**
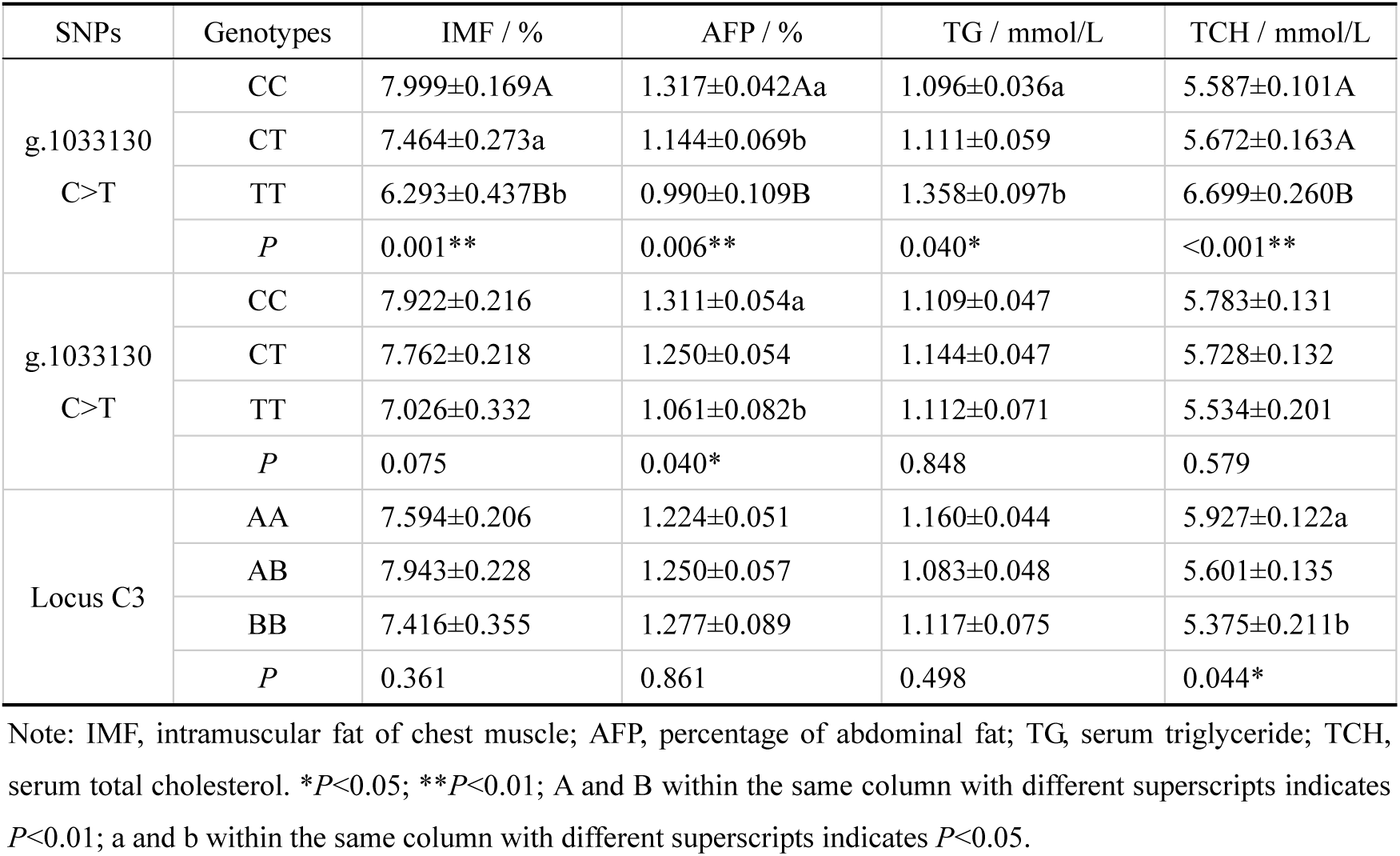
Association analysis of SNPs of CYP7A1 gene with lipid traits in Cherry Valley ducks.

### Associations between Diplotypes and Four Lipid Traits

The results of the association analysis between the diplotypes of haplotype combination and the lipid traits in Cherry Valley ducks were shown in Table 6. Our research indicated that the diplotypes had strongly associated with IMF, AFP, TG, and TCH respectively. The IMF of H1H1, H2H3, and H1H6/H4H5 diplotypes had greater than H5H6, H3H6, and H5H5 diplotype (*P*<0.01), and H1H6/H4H5 was the highest; the AFP of H2H3, H1H1, and H2H2 diplotypes had greater than H1H3 and H2H5 diplotypes (*P*<0.01), and H2H2 was the highest; the TG of H5H5 diplotype had greater than H1H1, H1H5, H1H6/H4H5, H2H3, H3H6, and H2H6 diplotypes (*P*<0.01), H1H3 and H3H5 diplotypes had greater than H1H1, H2H3, H3H6, and H2H6 diplotypes (*P*<0.01); the TCH of H5H6 diplotype had greater than other diplotypes except H5H5, H1H3 and H3H5 diplotypes (*P*<0.01), H5H5 diplotype had greater than H1H1, H1H5, H3H3, H2H5, H2H3, H1H6/H4H5, H2H6, H4H4, and H6H6 diplotypes (*P*<0.01), and H1H3 diplotype had greater than H2H3 and H4H4 diplotypes. In addition, there were significant or no significant difference with the lipid indexes between other diplotypes (*P*<0.05 or >0.05).

**Table 6.** Least square means and P-values of the effects of diplotypes on the lipid traits in Cherry Valley ducks.

| Diplotypes | IMF / % | AFY / % | TG / mmol/L | TCH / mmol/L |
| --- | --- | --- | --- | --- |
| H1H1 | 8.212±0.289Aab | 1.410±0.072Aa | 0.992±0.062Cc | 5.650±0.167CDcde |
| H1H5 | 8.001±0.640abcd | 1.261±0.158ab | 1.020±0.136BCc | 5.596±0.369CDcde |
| H5H5 | 6.241±0.678Bcd | 1.010±0.168b | 1.603±0.154Aa | 7.020±0.392ABab |
| H1H3 | 6.771±0.532bcd | 0.891±0.132Bb | 1.382±0.113ABab | 6.452±0.307ABCbc |
| H3H5 | 6.753±0.640bcd | 1.204±0.158ab | 1.460±0.136ABab | 6.257±0.369ABCDbcd |
| H3H3 | 7.449±0.554abcd | 1.031±0.137b | 1.160±0.118bc | 5.647±0.320CDcde |
| H1H2 | 7.968±0.640abcd | 1.299±0.158ab | 1.303±0.136abc | 5.942±0.369BCDcde |
| H2H5 | 7.661±0.725abcd | 0.884±0.180Bb | 1.111±0.154bc | 5.339±0.419CDde |
| H2H3 | 8.497±0.311Aa | 1.421±0.077Aa | 1.019±0.066Cc | 5.213±0.180De |
| H1H6/H4H5 | 8.611±0.607Aa | 1.190±0.150ab | 1.035±0.129BCc | 5.406±0.350CDde |
| H5H6 | 5.948±0.783Bd | 1.032±0.194ab | 1.305±0.167abc | 7.628±0.452Aa |
| H3H6 | 6.198±0.678Bd | 0.944±0.168b | 1.009±0.144Cc | 6.011±0.392BCDbcde |
| H2H2 | 8.103±0.678abc | 1.510±0.168Aa | 1.197±0.144abc | 5.699±0.392BCDcde |
| H2H6 | 7.318±0.678abcd | 1.314±0.168ab | 1.016±0.144Cc | 5.383±0.392CDde |
| H4H4 | 7.370±0.607abcd | 1.273±0.150ab | 1.127±0.129bc | 5.131±0.350De |
| H6H6 | 6.708±0.783bcd | 0.922±0.194b | 1.127±0.167bc | 5.342±0.452CDde |
| *P* | 0.005** | 0.003** | 0.007** | <0.001** |

Note: IMF, intramuscular fat of chest muscle; AFY, percentage of abdominal fat; TG, serum triglyceride; TCH, serum total cholesterol. \* $P<0.05$ ; \*\* $P<0.01$ ; A, B, C, D, E within the same column with different superscripts indicates $P<0.01$ ; a, b, c, d, e within the same column with different superscripts indicates $P<0.05$ .

## DISCUSSION

CYP7A1 emerges to play an important role in hypercholesterolemia and liver fat deposition, and apposite position of CYP7A1 has been targeted for innovative pharmacological intervention in recent years. Therefore, CYP7A1 was considered as a major candidate gene for lipid deposition in animal breeding. The previous studies reported that the polymorphisms of CYP7A1 gene had significantly associated with lipid indexes TG, TCH, LDL, and VLDL in human and mammal. The A-203C variant in promoter of CYP7A1 gene had significantly associated with plasma TG and low-density lipoprotein cholesterol (LDL-C) levels in different types of dietary treatment (Hubacek and Bobkova 2006; Vlachova *et al*. 2016). The combination of CYP7A1 rs3808607, DHCR7 rs760241 and ABCG5 rs6720173 is related with serum LDL-C concentrations to dairy consumption (Abdullah *et al*. 2018). CYP7A rs 2081687 is strongly associated with serum concentrations of TCH and LDL in different cohort characteristics (Teslovich *et al*. 2010). In avian species, however, little is known about the role of the polymorphisms of CYP7A1 gene except for the fact that it was involved in cholesterol catabolism and bile acid synthesis (Zhao *et al*. 2011; Sato *et al*. 2003; Chen *et al*. 2012).

In our study, we identified the seven novel synonymous mutations loci of CYP7A1 gene that showed significant association with at least one of the four lipid traits tested. Interesting five loci g.1034334G>A, g.1034373G>A, g.1034448T>C, g.1034541C>G, and g.1034550G>A were in complete linkage disequilibrium, and as a block named Locus C3. The g.1033130C>T locus had notable affected on lipid traits than g.1034076 C>T locus and Locus C3, and revealed that the function of DNA stranded conformational change of CYP7A1 gene was caused by g.1033130C>T locus may greater than g.1034076 C>T locus and Locus C3 for four lipid indexes, because previous many studies demonstrated that DNA conformational change may be effect on animal performance (Stefl *et al*. 2013). Totally six haplotypes and sixteen dipoltypes were found using SHEsis online analysis software, and showed significant difference between different diplotypes, suggested dominance/over-dominance effect on four lipid indicators between different haplotypes according to dominance/over-dominance hypothesis (Xu *et al*. 2013; Dekkers and Chakraborty 2004). Therefore, our results advised breeders to breed different strains base on the effect on various lipid indexes between different haplotypes or genotypes of each SNP in order to obtain heterosis.

Different haplotypes may produce different molecular conformation of DNA or RNA that influence the gene copy number, and the activity of the CYP7A1 promoter contains several regulatory elements such as liver X receptor (LXR) response element (LXRE), hepatocyte nuclear factor 4α (HNF-4α) binding site, and liver receptor homolog-1 (LRH-1) responsive element, and finally impact on the levels of transcription and translation via regulating the binding efficiency of transcription factors (such as LXRs et al) or ligands (such as FXR e al) or transcriptional repressor (small heterodimer partner, SHP) to CYP7A1 (Moon *et al*. 2016). Acetyl-coenzyme A (AcCoA) acts as an essential precursor molecule for both cholesterol and fatty acid synthesis. Transcription factors, LXRs and SREBPs, are considered the most major regulatory elements in fatty acid or cholesterol synthesis (Gbaguidi and Agellon 2004). SREBPs are critical transcription regulators of multiple genes linked in lipid metabolism. SREBP-1a and SREBP-1c mainly facilitated fatty acid biosynthesis, while SREBP2 is considered to be essential for cholesterol uptake through LDL receptors and de novo cholesterol biosynthesis through mevalonate pathway (Song *et al*. 2015). As sterol sensors, LXRs are responses to cellular sterol loading, forming obligate heterodimers with retinoid X receptors (RXRs) and triggering the expression of SREBP-1c and cholesterol efflux-related genes, and then induces triglyceride accumulation through fatty acid synthase (FAS) (Cai *et al*. 2016). Meanwhile, cholesterol is one component of serum LDL, and is transported into liver through LDL receptor, and induces the expression of CYP7A1 and cholic acid biosynthesis (Saneyasu *et al*. 2013). CYP7A1 as a target gene of LXRs, its low-level expression causes cholesterol catabolism disorder and cholic acid synthesis reduction. Based on the above research results, CYP7A1 signaling has been mentioned to participate in lipid metabolism, and a variety of molecules including LXRα, SREBP-1c, FXR, FAS, 3-hydroxy-3-methylglutaryl coenzyme A reductase (HMGR), lecithin cholesterol acyltransferase (LCAT), and the scavenger receptor CD36 were involved, and then regulate the dynamic balance of serum TC and TG concentrations through fatty deposition (body fatty and IMF) and oxidative decomposition (Chiang 2003; van Solingen *et al*. 2018; Zhang *et al*. 2018). Thus, different haplotypes may cause the differentiating expression of CYP7A1 lead to the differential expression of genes involved in lipid metabolism, and all of these genes together affect intramuscular fat (IMF) content of chest muscles, percentage of abdominal fat (AFP), serum triglycerides (TG) and total cholesterol (TCH) concentrations. Certainly, further work is necessary, such as the effects of each SNP and haplotypes/diplotypes on expression of CYP7A1 will be evaluated in different duck populations, and the effect of CYP7A1 gene expression on lipid metabolism-related genes should be confirmed in the future.

## CONCLUSIONS

In our research, seven novel synonymous mutations loci in exon 2 and exon3 of CYP7A1 gene were identified in Cherry Valley ducks, and each SNP showed significant association with at least one of the four lipid traits tested. Six haplotypes were detected. Sixteen dipoltypes were found by the combination of haplotypes, and showed significant difference between different diplotypes for four lipid indexe, intramuscular fat (IMF) content of chest muscles, percentage of abdominal fat (AFP), serum triglycerides (TG) and total cholesterol (TCH) concentrations. Therefore, the seven SNPs of CYP7A1 were potential markers for lipid metabolism balance. These findings may be instructional for early breeding and selection of duck.

## AUTHOR CONTRIBUTIONS

Yuan-Yu Qin performed the experiments, analyzed the data, prepared the figures and tables, and prepared the manuscript.

Yi-Yu Zhang is project leader, conceived and designed the study, and revised the manuscript.

Hua-Lun Luo attended the experimental operation and analyzed the data of the study. Lei Wu reared the experimental animals and measured the data of the study.

## ACKNOWLEDGMENTS

This research was financially supported by the National Natural Science Foundation of China (31760663) and Science Technology Project of Guizhou Province of China (QKHPTRC[2017]5788).

## ETHICAL STATEMENT

All subjects gave their informed consent for inclusion before they participated in the study. All animal experiments were performed according to the Laboratory animal—Guideline for ethical review of animal welfare of China (permit number: GB/T 35892-2018) that was issued by China Laboratory Animal Standardization Technology Committee (SAC/TC 281). All procedures were performed under anesthesia and euthanasia. Animals were closely monitored and observed for development of disease once daily, and all efforts were made to minimize suffering.

